# Immunological features that determine the strength of antibody responses to BNT162b2 mRNA vaccine against SARS-CoV-2

**DOI:** 10.1101/2021.06.21.449182

**Authors:** Takahiro Kageyama, Shigeru Tanaka, Keishi Etori, Koto Hattori, Kazusa Miyachi, Tadamichi Kasuya, Taro Iwamoto, Kei Ikeda, Hidetoshi Igari, Koutaro Yokote, Hiroshi Nakajima

## Abstract

We analyzed peripheral blood mononuclear cells (PBMCs) of each 20 individuals with a high anti-SARS-CoV-2 antibody titer and a low antibody titer out of 1,774 healthcare workers who received BNT162b2 mRNA vaccine. A higher antibody titer was associated with the frequencies of naïve and transitional B cells before vaccination. In addition, fold changes in the frequency of activated CD8^+^ T cells upon vaccination were correlated with the antibody titers.

## MAIN TEXT

It has been reported that the BNT162b2 mRNA vaccine contributed to reducing the severity of COVID-19 (1). Vaccination against SARS-CoV-2 is progressing around the world at an unprecedented rate (2). However, the pandemic of COVID-19 has led to many SARS-CoV-2 variants, some of which have been highly transmissible and partially resistant to immune responses obtained from previous infection or vaccination (3). Although BNT162b2 has been shown to induce vaccine-elicited neutralization against SARS-CoV-2 variants so far (4, 5), it may be required to improve vaccines before the virus acquires critical mutations.

As the humoral responses play vital roles in the protection against SARS-CoV-2 infection (6, 7), the antibody titer status after vaccination can provide essential information to develop better vaccines and optimize vaccination strategies. We have previously reported favorable antibody responses to BNT162b2 and their predictive clinical factors in the largest study to date (8).

Although age has been repeatedly shown to be associated with a lower antibody response among demographic factors (9,10), there is a subgroup with low antibody titers even in young populations without well-known factors reducing antibody responses, such as taking immunosuppressive agents and glucocorticoids (8). Therefore, we aimed to clarify the immunological backgrounds that underlie the difference in antibody responses. To address this issue, we investigated immunophenotypic characteristics in peripheral blood mononuclear cells (PBMCs), collected both before and after vaccination, among high and low responders to the BNT162b2 mRNA COVID-19 vaccine.

We selected 20 young individuals with a low antibody titer after vaccination out of 1,774 healthcare workers who received 2 doses of BNT162N2 and 20 age- and sex-matched individuals with a high antibody titer. Individuals with a history of COVID-19 or a detectable antibody titer (≥0.4 U/mL) before the vaccination were excluded from the study. Individuals who were taking oral glucocorticoids or immunosuppressive medication were also excluded. The median antibody titer was 4,965 U/mL (interquartile range [IQR] 3,888-6,530) in the high responder group, while the median antibody titer was 445.0 U/mL (IQR 325.5-552.3) in the low responder group. Detailed background information is provided in Table 1.

**Table 1.** Background information of high and low responders to the BNT162b2 vaccine

|  | High responders (n = 20) | Low responders (n = 20) |
| --- | --- | --- |
| Age, year, median (IQR) | 33.5 (28.0-40.5) | 35.5 (26.8-40.3) |
| Sex (female:male), n | 10:10 | 9:11 |
| Post-vaccine anti-SARS-CoV-2S antibody titer, U/mL, median (IQR) | 4,965 (3,888-6,530) | 445.0 (325.5-552.3) |
| Alcohol consumption |  |  |
| No, n (%) | 8 (40) | 4 (20) |
| Sometimes, n (%) | 12 (60) | 13 (65) |
| Every day, n (%) | 0 (0) | 3 (15) |
| Smoking |  |  |
| Never, n (%) | 17 (85) | 15 (75) |
| Ex-smoker, n (%) | 2 (10) | 3 (15) |
| Current smoker, n (%) | 1 (5) | 2 (10) |
| Time between 1<sup>st</sup> and 2<sup>nd</sup> dose, day, mean (SD) | 21.1 (0.5) | 21.0 (0.2) |
| Time between 2<sup>nd</sup> dose and sample collection, day, median (IQR) | 14 (14-21) | 15 (14-21) |
| Comorbidities |  |  |
| Hypertension, n (%) | 1 (5) | 1 (5) |
| Dyslipidemia, n (%) | 1 (5) | 1 (5) |
| Diabetes mellitus, n (%) | 0 (0) | 1 (5) |
| Malignancy, n (%) | 1 (5) | 0 (0) |
| Atopic dermatitis, n (%) | 2 (10) | 1 (5) |
| Asthma, n (%) | 7 (35) | 2 (10) |
| Thyroid disease, n (%) | 0 (0) | 1 (0) |
| Current medication |  |  |
| Hypertension, n (%) | 1 (5) | 1 (5) |
| Dyslipidemia, n (%) | 1 (5) | 0 (0) |
| Diabetes mellitus, n (%) | 0 (0) | 1 (5) |
| Allergy, n (%) | 2 (10) | 1 (5) |
| Inhaled corticosteroid, n (%) | 2 (10) | 0 (0) |
| Glucocorticoids, n (%) | 0 (0) | 0 (0) |
| Immunosuppressant, n (%) | 0 (0) | 0 (0) |
| Thyroid disease, n (%) | 0 (0) | 1 (5) |

IQR, interquartile range. SD, standard deviation.

We quantified the frequencies of CD4^+^ and CD8^+^ T cells, B cells, and monocytes in PBMCs, as shown in Fig S1. Interestingly, the percentage of B cells was positively correlated with a higher antibody titer, while the percentages of T cells and monocytes were not (Fig 1a). Among B cells, naïve and transitional B cell frequencies were positively correlated with a higher antibody titer, whereas the frequencies of late memory B cells and plasmablasts were associated with a lower antibody titer (Fig 1a). To our surprise, the frequencies of CD4^+^ T cell subsets (e.g., naïve, effector, effector memory, and central memory) were not significantly associated with antibody titers despite the well-known roles of CD4^+^ T cells in antibody responses. Also, there was no correlation between antibody titers and intermediate monocytes, which are important for antigen presentation. These results suggest that the abundance of naïve and immature B cells is associated with the robust antibody response against the BNT162b2 vaccine.

**Figure 1.**
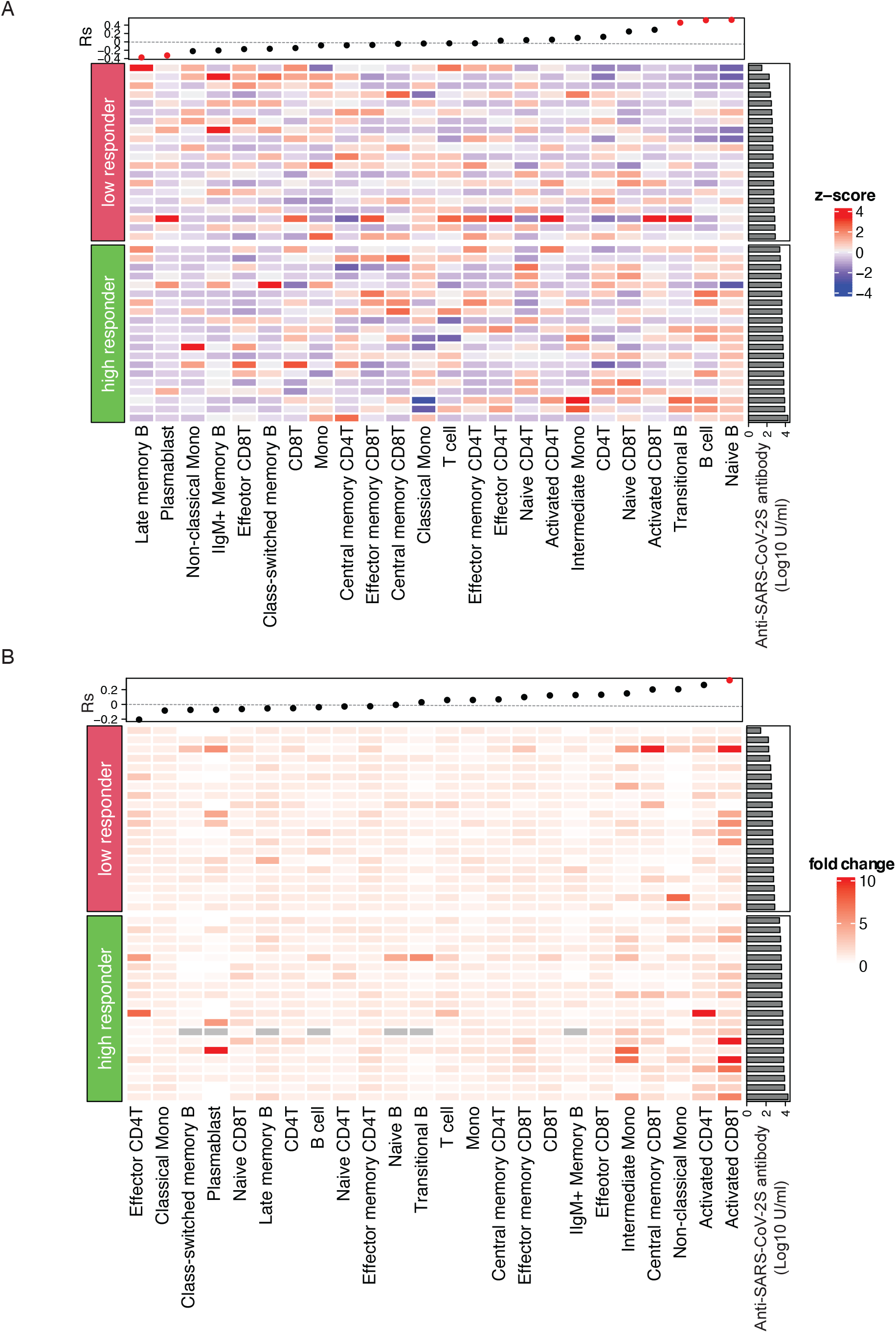
The correlation between the proportion of immune cell subsets in PBMCs and the anti-SARS-CoV-2S antibody titer. (**A, B**) One line of the heatmap and the bar diagram on the right represents the data of each individual. The heatmap shows the z-scored percentage of each cell subset in PBMCs before vaccination (**A**) and the fold change of each cell subset in PBMCs after vaccination (**B**). Seven gray tiles indicate missing data due to technical errors. The bar diagram displays anti-SARS-CoV-2S antibody titer (Log_10_U/mL) after vaccination. Spearman’s rank correlation coefficient Rs between anti-SARS-CoV-2S antibody titer and the percentage of each cell subset are plotted above the heatmap. Red dots indicate those with a significant difference.

We also analyzed the changes in cell fractions after the vaccination. The increased frequency of activated CD8^+^ T cells was positively correlated with a higher antibody titer (Fig 1b). In addition, there was a trend to have higher induction of activated CD4^+^ T cells in high responders (Fig 1b). These results suggest that the robust T cells responses on the BNT162b2 vaccine are associated with high antibody responses.

Recently, through the analysis of human naïve B cells, some naïve antibodies have been shown to bind to the receptor-binding domain of SARS-CoV-2 and can neutralize SARS-CoV-2 pseudoviruses even in the absence of somatic mutations (11,12). Furthermore, although the clinical settings are different from our study, kidney transplant recipients and dialysis patients have diminished humoral responses to BNT162b2 with reduced B cells (13). These results suggest that the abundance of naïve B cells may be associated with the presence of a diverse BCR repertoire, which induces a good antibody response. This notion is consistent with previous findings that high antibody titers are significantly associated with younger age groups (8, 9, 10), who are thought to have more naive B cells. Meanwhile, some studies have examined the relationship between B cell fractions and antibody production with influenza vaccines, the most widely used vaccine in humans, and have reported that naïve B cells do not correlate with high antibody responses (14, 15). We speculate that the discrepancy between the current study on the SARS-CoV-2 vaccine and studies on influenza vaccines may be due to the presence of virus-specific memory B cells by previous infections or vaccines (16). We also speculate that immune responses to the mRNA vaccine may differ from those to the usual inactivated vaccine. T cell activation after the vaccination is also associated with high antibody titers (Fig 1B), which may suggest broad immunogenicity of the mRNA vaccine. Further studies are required to know whether the relation between naïve B cells and the response to BNT162b2 is still observed if vaccination against COVID-19 is repeated over the years.

In conclusion, we have demonstrated that high proportions of naïve and transitional B cells before vaccination are associated with good responses to the BNT162b2 vaccine, although the mechanisms remain unknown. Since BNT162b2 is the first mRNA vaccine to be widely used in humans, this study has important insights into vaccine development. In order to control the pandemic of COVID-19, we need to promote faster and more effective vaccination, analyzing the data obtained from ongoing vaccinations.

## Figure Legends

**Figure S1.**
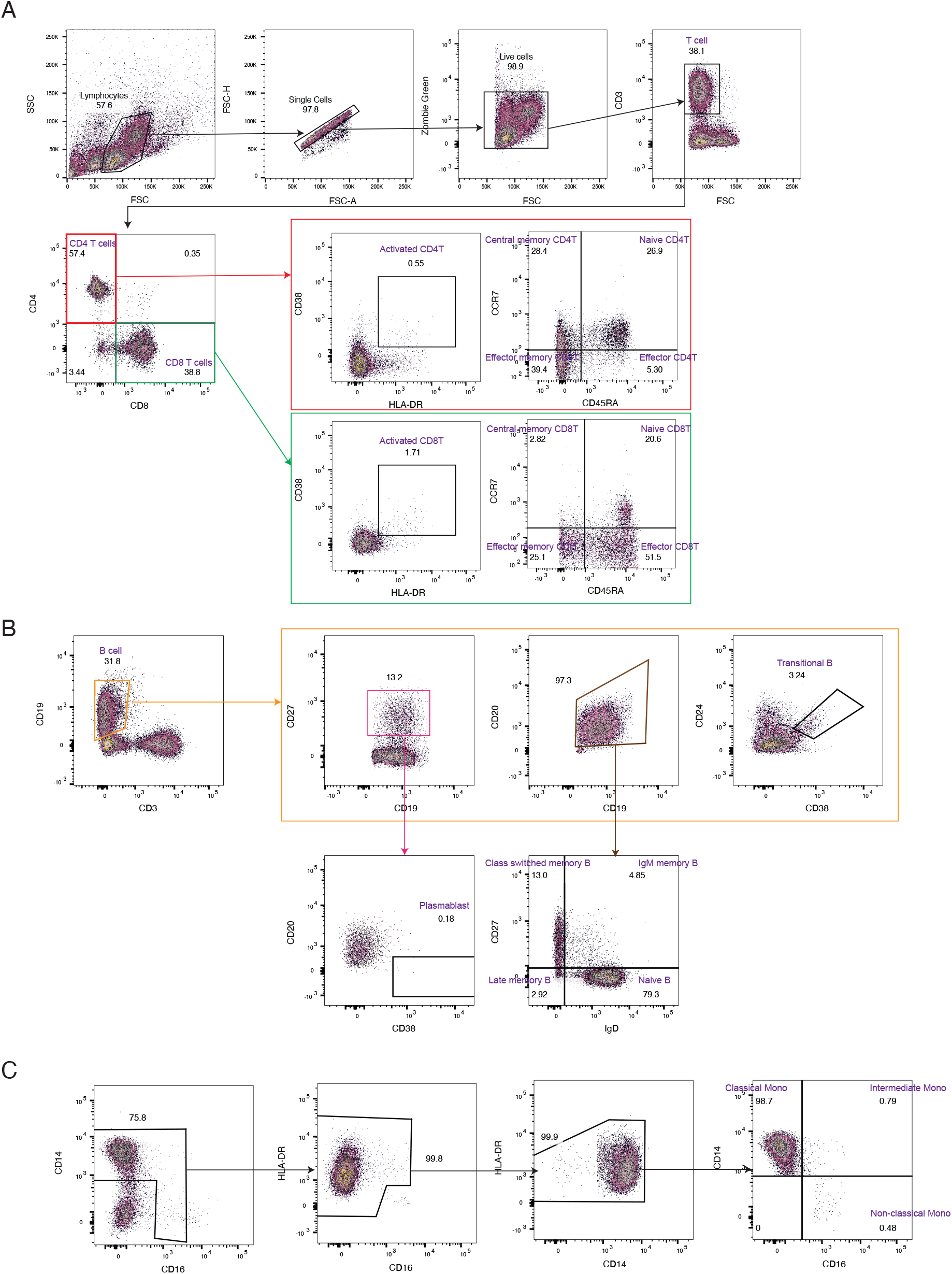
Representative pseudocolor plots showing the gating strategy for immunophenotypic characteristics in PBMCs. (**A**) Representative pseudocolor plots illustrate gating strategies for identifying activated, central memory, effector memory, effector, naïve CD4^+^ or CD8^+^ T cells. Activated CD4^+^ or CD8^+^ T cells were classified based on CD38 and HLA-DR. Central memory, effector memory, effector, and naïve CD4^+^ or CD8^+^ T cells were classified based on CCR7 and CD45RA. (**B**) Representative pseudocolor plots illustrate gating strategies for identifying B cells, plasmablasts, class-switched memory B cells, late memory B cells, IgM^+^ memory B cells, naïve B cells, and transitional B cells. Class-switched memory B cells, late memory B cells, IgM^+^ memory B cells, and naïve B cells were classified based on CD27 and IgD. (**C)** Representative pseudocolor plots illustrate gating strategies for identifying classical monocytes, intermediate monocytes, and non-classical monocytes. Classical, intermediate, and non-classical monocytes were classified based on CD14 and CD16.

## METHODS

### Participants

We recruited healthcare workers in Chiba University Hospital who received the BNT162b2 mRNA COVID-19 vaccine (Pfizer, Inc., and BioNTech) at Chiba University Hospital COVID-19 Vaccine Center (8). Among them, we selected 20 high responders and 20 low responders against the BNT162b2 mRNA vaccine based on anti-SARS-CoV-2S antibody titer. They were between their 20s – 40s in age and did not have a history of COVID-19. No one takes oral steroids or immunosuppressive agents. The study procedures for sample collection and those for analyses were approved by Chiba University Ethics Committee on February 24th, 2021 (No. HS202101-03) and April 21st, 2021 (No. HS202104-01), respectively.

### Sample collection and peripheral blood mononuclear cell preparation

Blood samples were obtained 0-2 weeks before the 1st dose and 3 weeks after the 2nd dose of vaccination. Peripheral blood mononuclear cells (PBMCs) were stored in liquid nitrogen until analysis.

### Anti-SARS-CoV-2S antibody test

Anti-SARS-CoV-2S antibody was measured by Elecsys® Anti-SARS-CoV-2S on Cobas 8000 e801 module as described elsewhere (8).

### Flow cytometry analyses

PBMCs were first stained with either Zombie Green (for T cell and B cell staining panel) (Biolegend) or Zombie NIR (for monocyte staining panel) (Biolegend) to label dead cells. Then the samples were treated with Human TruStain FcX (Biolegend) to block Fc receptors. For the analysis of T cells, cells were stained with antibodies against CCR7 (150503, BD), CD4 (RPA-T4, Biolegend), CD45RA (HI100, BD), CD38 (HIT2, BD), CD8 (SK1, BD), CD3 (UCHT1, BD), and HLA-DR (G46-6, BD). For the analysis of B cells, cells were stained with antibodies against CD24 (ML5, BD), CD19 (SJ25C1, BD), CD27 (O323, Biolegend), CD38, CD20 (2H7, BD), and IgD (IA6-2, BD). For the analysis of monocytes, cells were stained with antibodies against CD16 (B73.1, BD), CD14 (MfP9, BD), and HLA-DR. The gating strategies were shown in Figure S1. Data were collected by a Canto II (BD) and analyzed by FlowJo software (BD).

### Analyses and visualization

The percentages of each cell subset were evaluated for each individual. T cells, B cells, and monocytes are shown as % in live cells. CD4^+^ T cells and CD8^+^ T cells are shown as % in T cells. Activated CD4^+^ T cells, naïve CD4^+^ T cells, effector CD4^+^ T cells, central memory CD4^+^ T cells, and effector memory CD4^+^ T cells are shown as % in CD4^+^ T cells. Activated CD8^+^ T cells, naïve CD8^+^ T cells, effector CD8^+^ T cells, central memory CD8^+^ T cells, and effector memory CD8^+^ T cells are shown as % in CD8^+^ T cells. Plasmablasts, naïve B cells, IgM^+^ memory B cells, class-switched memory B cells, late memory B cells, and transitional B cells are shown as % in B cells. Classical monocytes, intermediate monocytes, and non-classical monocytes are shown as % in monocytes. Spearman’s rank correlation test was used to analyze the correlation between anti-SARS-CoV-2S antibody titer and the percentage of each cell subset. For Figure 1A, the heatmap shows the z-scored percentage of each cell subset along with the antibody titer and Spearman’s rank correlation coefficient Rs. For Figure 1B, the fold change of each cell subset after the vaccination was calculated, and the correlation with the antibody titer was tested using Spearman’s rank correlation coefficient. P values less than 0.05 were considered statistically significant.

## DATA AVAILABILITY

All data generated or analyzed during this study are included in this published article and its supplementary information files.

## ACKNOWLEDGEMENTS

We thank all staff in Chiba University Hospital for supporting sample collection. This study was supported by a grant from JST [Moonshot R&D] [Grant Number JPMJMS2025] and a donation to Chiba University Hospital and the Future Medicine Foundation at Chiba University.

## AUTHOR CONTRIBUTIONS

T.K., S.T., and H.N. designed the study; T.K., S.T., K.E., K.M., and T.K. performed the flow cytometry analysis; T.K., S.T., T.I., K.I., H.I, K.Y, and H.N. interpreted the data; T.K., S.T., and H.N. wrote the manuscript.

